# A Data-Driven Latent Variable Approach to Validating the Research Domain Criteria Framework

**DOI:** 10.1101/2024.01.31.577486

**Authors:** S.K.L. Quah, B. Jo, C. Geniesse, L.Q. Uddin, J.A. Mumford, D.M. Barch, D.A. Fair, I.H. Gotlib, R.A. Poldrack, M. Saggar

## Abstract

Despite the widespread use of the Research Domain Criteria (RDoC) framework in psychiatry and neuroscience, recent studies suggest that the RDoC is insufficiently specific or excessively broad relative to the underlying brain circuitry it seeks to elucidate. To address these concerns, we employed a latent variable approach using bifactor analysis. We examined 84 whole-brain task-based fMRI (tfMRI) activation maps from 19 studies with 6,192 participants. A curated subset of 37 maps with a balanced representation of RDoC domains constituted the training set, and the remaining held-out maps formed the internal validation set. External validation was conducted using 36 peak coordinate activation maps from Neurosynth, using terms of RDoC constructs as seeds for topic meta-analysis. Here, we show that a bifactor model incorporating a task-general domain and splitting the cognitive systems domain better fits the examined corpus of tfMRI data than the current RDoC framework. We also identify the domain of arousal and regulatory systems as underrepresented. Our data-driven validation supports revising the RDoC framework to reflect underlying brain circuitry more accurately.

## Introduction

The study of human neurobiology is a rapidly advancing field with significant implications for understanding brain function and, eventually, facilitating the development of valid biological markers and effective treatments for psychiatric disorders. Psychiatric disorders listed in the Diagnostic and Statistical Manual (DSM) have historically been considered to be discrete and unitary; recent research, however, suggests that they are both highly comorbid and heterogeneous across clinical samples^1,2^. This heterogeneity may underlie the lack of well-established biomarkers to date for psychiatric disorders.

The Research Domain Criteria (RDoC) framework was developed by the National Institute of Mental Health (NIMH) to guide the development of a psychiatric nosology based on primary psychological functions and their associated biological features^3,4^. The framework organizes core dimensions of behavior using a dimensional approach, viewing these aspects as varying along a continuum rather than in distinct categories. This approach spans multiple levels of analysis, from genes to behavior^5^. Within the RDoC framework, the fundamental neurobiological systems were defined and organized hierarchically into domains, with domain-specific constructs and sub-constructs. Now, over a decade since its inception, the framework’s dimensional approach to psychopathology and its integration of multiple levels of analysis have contributed to a more nuanced and comprehensive understanding of brain function and mental disorders^4,6^.

While the RDoC framework has helped guide research, a recent study using text-mining and machine learning found that a bottom-up data-driven ontological framework generated brain circuit-function links that were more reproducible than the RDoC or DSM frameworks^7^. They also showed that multiple RDoC domains shared underlying neural circuits or some domains needed to be split. For example, Beam et al.^7^ showed that the RDoC domains of negative valence, positive valence, and arousal and regulatory systems shared high mutual information across the fronto-medial cortex and amygdala, indicating an overlap in the division of these domains. Further, they also showed that the RDoC negative valence domain encompassed constructs that, from a data-driven framework, recombine elements of memory, reward, and cognitive systems. These findings prompt further investigation into potential refinements to RDoC’s domain structure and mapping of brain function to neural circuits.

Researchers have made significant strides in attempting to develop a data-driven ontology that maps brain function to neural circuits through the meta-analysis of task-based fMRI (tfMRI) activation maps and topic modeling. Using data mining techniques, peak brain coordinate activation patterns during tasks have been categorized based on latent functional domains derived from study texts^8,9^ or task descriptions^10,11^. While previous studies utilizing coordinate activation data have effectively harnessed the vast amounts of data available in databases like Neurosynth^12^ and Brainmap^13^, they provide a very sparse representation of whole-brain activation. Image-based meta-analyses can provide a richer understanding of the intricate patterns of activation that occur during tasks^14^. It would be beneficial to compare RDoC directly with a data-driven model derived using image-based analyses to assess potential refinements to its framework.

To expand on the RDoC framework’s hierarchical structure and address any potential overlap between domains or lack of specificity within a domain, we leveraged a latent variable approach with bifactor analysis to explore circuit-function relations. Bifactor models allow one to capture both shared variance across a number of latent constructs as well as variance unique to specific constructs. Assessing both general patterns of brain activity common across tasks^15,16^ and task-specific activation, Bolt et al. previously demonstrated that a bifactor model represents the relations between psychological constructs and underlying neural processes better than conventional non-hierarchical frameworks^17^. Using a bifactor model can help to identify shared and unique variance among the different constructs and provide more nuanced insight into the organization of circuit-function relations. This approach can also help identify constructs that may be better conceptualized as part of a larger domain rather than as separate constructs. In this context, we used a bifactor analysis to examine the hierarchical structure of the RDoC framework across domains to provide data-driven evidence of complementary domain structures.

Specifically, we applied a latent variable approach with bifactor analysis to whole-brain task activation images from Neurovault and U.K. Biobank (n=84 select activation maps from 19 studies with a total of N=6,192 participants; adapted from Bolt et al^17^) to examine the organization of circuit-function relations. To ensure the robustness of our findings, we first derived our model solutions via a curated subset of the original dataset. Subsequently, we tested the model solution by applying it to the held-out maps, assessing its ability to generalize to previously unseen data. Moreover, we validated further using maps reconstructed from activation coordinates sourced from Neurosynth to assess the model’s applicability to diverse data types. This comprehensive approach (Fig. 1) allows us to evaluate how well our model solution captures and represents brain activation patterns across various datasets and serves as a crucial step in advancing our understanding of circuit-function relations.

**Figure 1:**
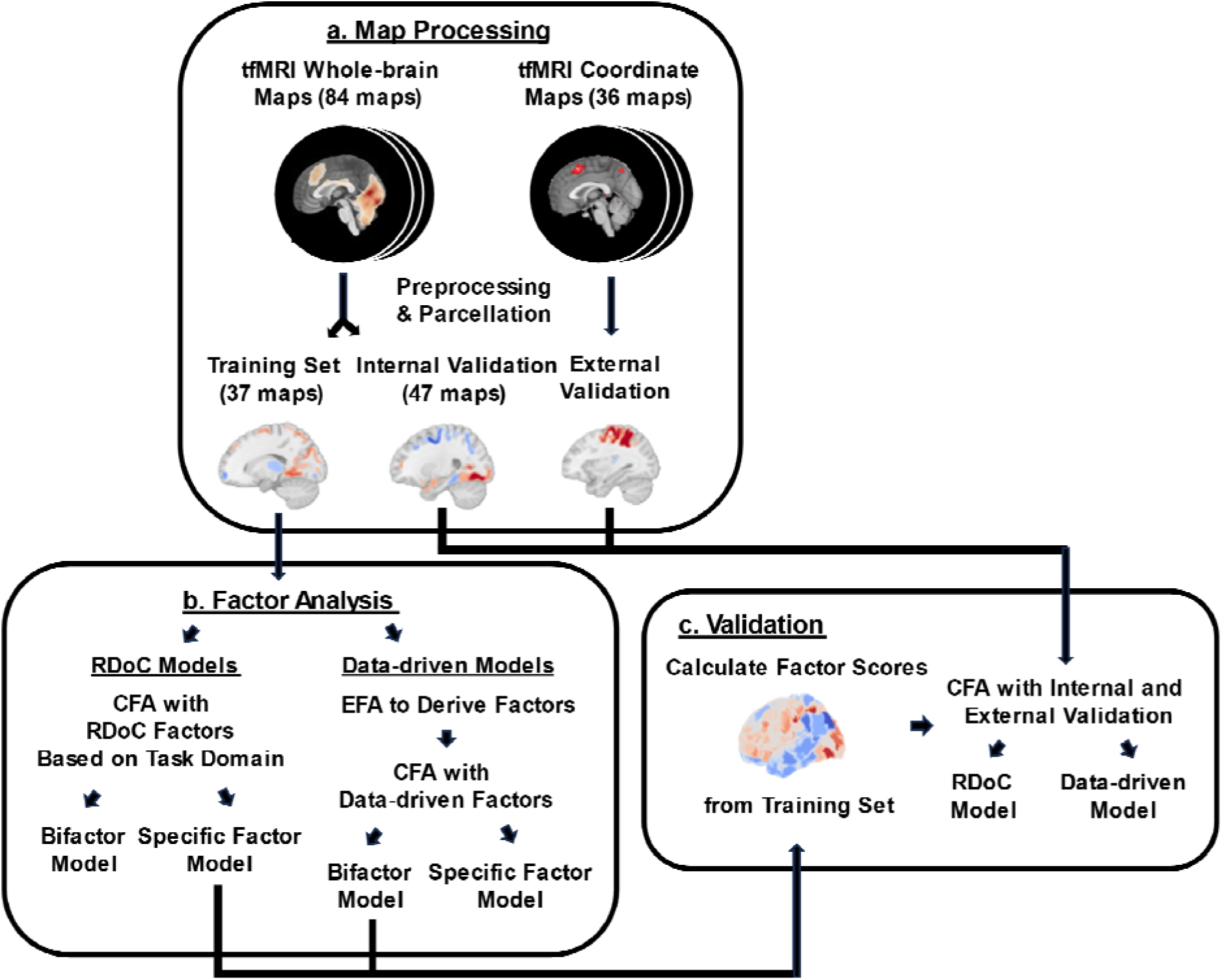
Approach to create and validate RDoC and data-driven factor models. (a) First, we divided tfMRI whole-brain activation maps into two subsets: the curated training dataset for building our factor models and the other as the internal validation set. Additionally, we processed tfMRI coordinate activation maps of peak activations to create an external validation set. (b) Regarding the confirmatory factor analysis (CFA) for the RDoC factor models, we assigned maps from the curated training dataset to specific factors corresponding to RDoC domains based on task associations. Before conducting a CFA, we first performed an exploratory factor analysis (EFA) to determine factor assignments for data-driven factor models. (c) We employed a validation procedure to evaluate the model’s performance on unseen data. We assigned maps to specific factors based on factor scores derived from the original data-driven and RDoC models. We then compared fit scores to assess the model’s generalizability to new data.

In this work, we demonstrate that a bifactor model, incorporating a task-general domain and refining the cognitive systems domain, provides a better fit to task-based fMRI data than the current RDoC framework. Our results suggest that refinements to the RDoC framework, informed by data-driven insights, could better capture the complexity of brain circuitry.

## Results

Latent variable models are designed to estimate latent constructs or classes that are not observed directly but are inferred from observed variables with measurement error^18^. We conducted a comparative analysis of four distinct latent variable approaches, combining two methods of factor derivation (theory-driven RDoC factors or data-driven empirical factors) with two types of factor models (specific factor models or bifactor models). Specific factor models exclusively incorporate specific factors, while bifactor models have an additional general factor^19^. To summarize, our study compared four models with the curated training dataset: (i) an RDoC specific factor model; (ii) an RDoC bifactor model; (iii) a data-driven specific factor model; and (iv) a data-driven bifactor model (Fig. 2).

**Figure 2:**
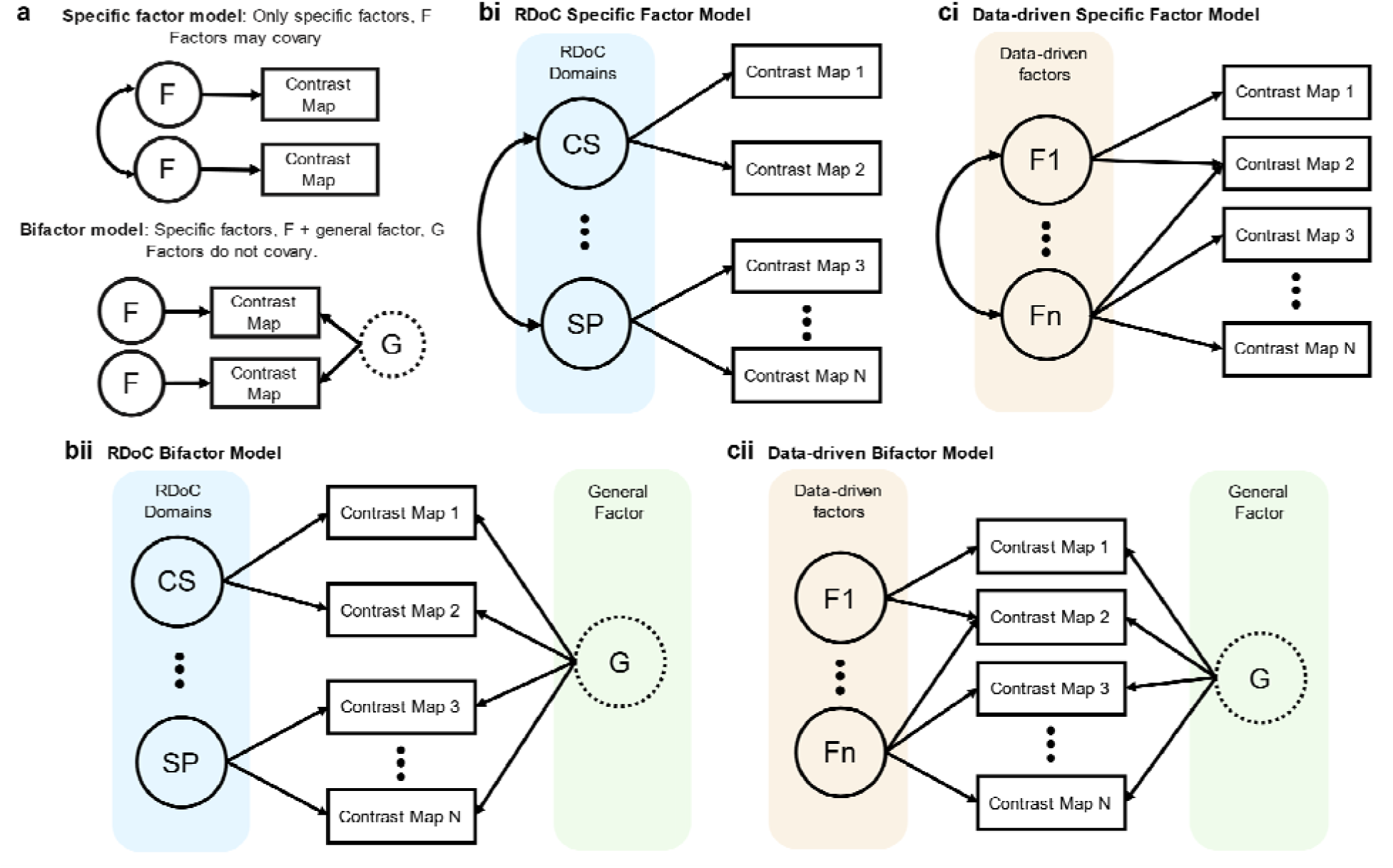
Factor model types. (a) Across all models, specific factors, F, denote brain activation patterns unique to a subset of tasks. In the bifactor models, the general factor, G, embodies brain activation patterns common across various tasks. (bi-bii) RDoC Models: These models are characterized by specific factors, each representing a distinct RDoC domain defined by task contrasts from whole-brain or coordinate activation maps. The Specific Factor Model (bi): This model comprises specific factors. The Bifactor Model (bii): This model is an extension of the specific factor model, with the addition of a general factor. (ci-cii) Data-Driven Models: These models (i & ii) are generated through EFA without predefined factors. CS and SP are representative RDoC domains, Cognitive Systems, and Social Processes, respectively.

### RDoC models with whole-brain activation maps

We conducted two CFAs with RDoC factors: one with only specific factors (Fig. 2bi) and another with an additional general factor (bifactor model; Fig. 2bii). Based on the task description of each contrast map (Supplementary Table 1), maps were grouped into specific factors by matching respective RDoC domains’ definitions.

In the specific factor model, most maps within each domain loaded significantly (i.e., |loading score|>=0.4) onto each factor representing their domains (cognitive systems: 11/15; negative valence systems: 5/5; positive valence systems: 6/7; social processes: 6/6; sensorimotor systems: 4/4; Fig. 3a).

**Figure 3:**
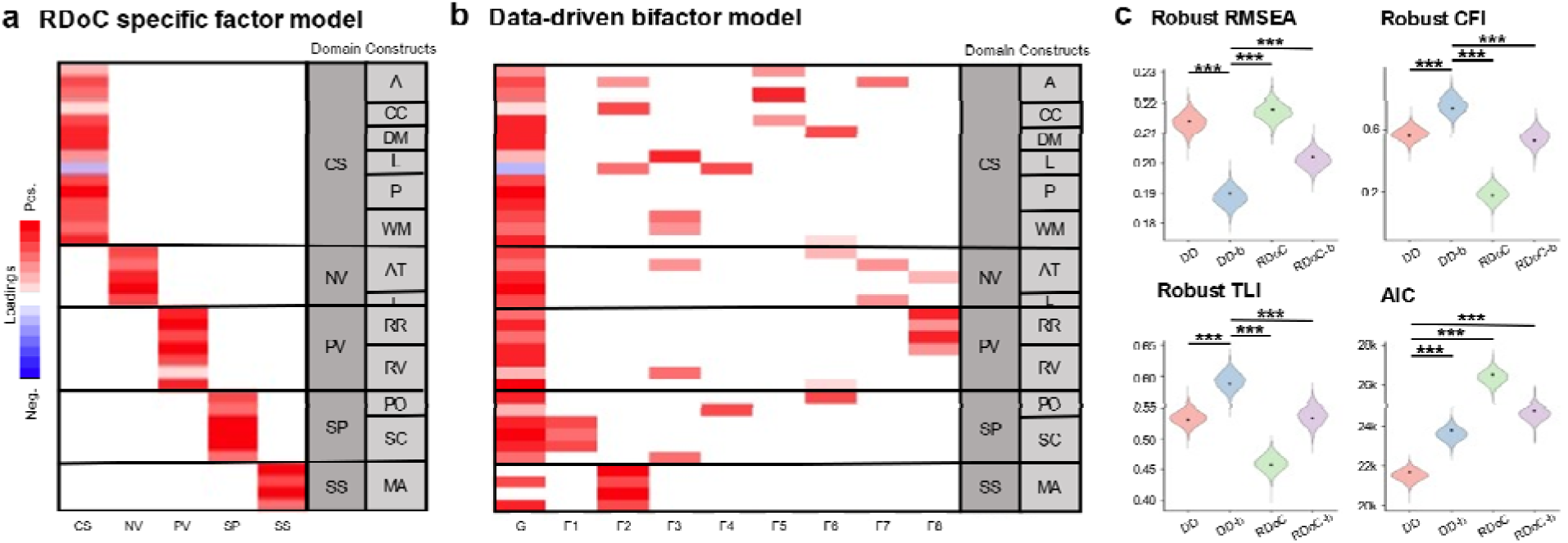
Comparison of RDoC and Data-driven models using whole-brain activation maps in the curated training set. (a-b) Heatmaps showing factor loadings of the RDoC and data-driven models across RDoC domain classified maps. Warmer colors = positive; Cooler colors = negative factor loadings shown. (c) Relative fit measures of different latent variable models from whole-brain activation maps. The data-driven bifactor model (DD-b) was the model with the best fit based on robust RMSEA, CFI, and TLI, but the data-driven specific factor model (DD) was the model with the best fit based on AIC. The violin plots represent the distribution of the bootstrap samples; the fit values from our models are indicated with black dots. CS: Cognitive Systems; NV: Negative Valence systems; PV: Positive Valence systems; SS: Sensorimotor Systems; SP: Social Processes; G: General factor; F1-8: specific factors 1-8; and DD: Data-driven. Domain-Constructs: Cognitive Systems-A: Attention; CC: Cognitive Control; DM: Declarative Memory; L: Language; P: Perception; WM: Working Memory; Negative Valence systems-AT: Acute Threat; L: Loss; Positive Valence Systems-RR: Reward Response; RV: Reward Valuation; Social Processes-PO: Perception of Others; SC: Social Communication; Sensorimotor Systems-MA: Motor Action. RDoC: RDoC specific factor model; RDoC-b: RDoC bifactor model.

Comparing the RDoC specific factor model with the bifactor model to examine whether adding a general factor would improve the fit, we found that the bifactor model had a better fit according to all fit indices (Tukey’s test, *p* < .001). This suggests that adding a general factor reflecting domain-general activation patterns improved the model fit of the conventional RDoC framework. This was also true after accounting for model complexity (with the AIC and BIC score) in the additional number of parameters estimated in the bifactor model, indicating that adding a general factor also provided a better balance between fit and complexity.

### Data-driven models with whole-brain activation maps

In the data-driven approach, we also conducted two CFAs: one with only specific factors (Fig. 2ci) and another with an additional general factor (bifactor model; Fig. 2cii). The specific factors for both models are latent variables derived using EFA that account for the unique variance among subsets of activation maps. They represent dimensions of task activation patterns that are not shared across all maps. Parallel analysis was first conducted to determine the appropriate number of factors to extract from the dataset. The parallel analysis indicated that models with eight factors or less had eigenvalues greater than expected by chance (Supplementary Figure 1). Thus, we extracted eight specific factors in the data-driven CFAs.

In the data-driven bifactor CFAs, all maps loaded significantly (i.e., |loading score|>=0.4) on the general, specific, or both factors. All but two maps across RDoC domains loaded on the general factor, indicating that maps across distinct studies and tasks showed overlap in activation patterns (Fig. 3b). Notably, the two maps that did not load on the general factor were associated with contrasts related to button pressing in response to an auditory cue; in contrast, the tasks in the dataset primarily revolved around responses to visual cues.

Furthermore, maps labeled by RDoC domains showed divergent patterns in loadings across specific factors (Fig. 3b). Positive valence systems, social processes, and sensorimotor systems domain maps showed high loadings that were confined to relatively few specific factors. In contrast, cognitive and negative valence systems domain maps showed significant loadings spread across multiple specific factors.

The ANOVA results indicated significant differences in fit among all the RDoC and data-driven model types (robust RMSEA: *F*(3, 19588) = 108,961, *p* < .001; robust CFI: *F*(3, 19588) = 212,411, *p* < .001; robust TLI: *F*(3, 19588) = 209,379, *p* < .001; AIC: *F*(3, 19588) = 126,142, *p* < .001; BIC: *F*(3, 19588) = 87,435, *p* < .001). The data-driven bifactor model also had a greater overall fit to the data compared with both RDoC models and the data-driven specific factor model (Tukey’s test, *p* < .001; Fig. 3c). However, after accounting for the different number of parameters estimated in the models, the data-driven bifactor model had a better model fit than the RDoC specific factor model but not the data-driven specific factor model (Tukey’s test, *p* < .001; Fig. 3c). All Tukey pairwise comparisons are shown in Supplementary Table 2.

After deriving these models, we created a product matrix to study similarities in map loadings across factors from the RDoC specific factor model and the data-driven bifactor model (Fig. 4a). The values in the product matrix represent the average product of absolute non-zero value factor loadings in both models. The values range from 0-1, where 1 represents a complete 1-to-1 similarity in map loadings, and 0 represents no overlap. This matrix provides insight into the consistency of the boundaries within and without the RDoC domains. Maps of domains with cross-loading on many specific factors reflect heterogeneity within the domain’s boundaries (low intra-domain consistency); maps of domains that share high loading with other domains on the same specific factor reflect overlap in the domains’ boundaries (high inter-domain similarity).

**Figure 4:**
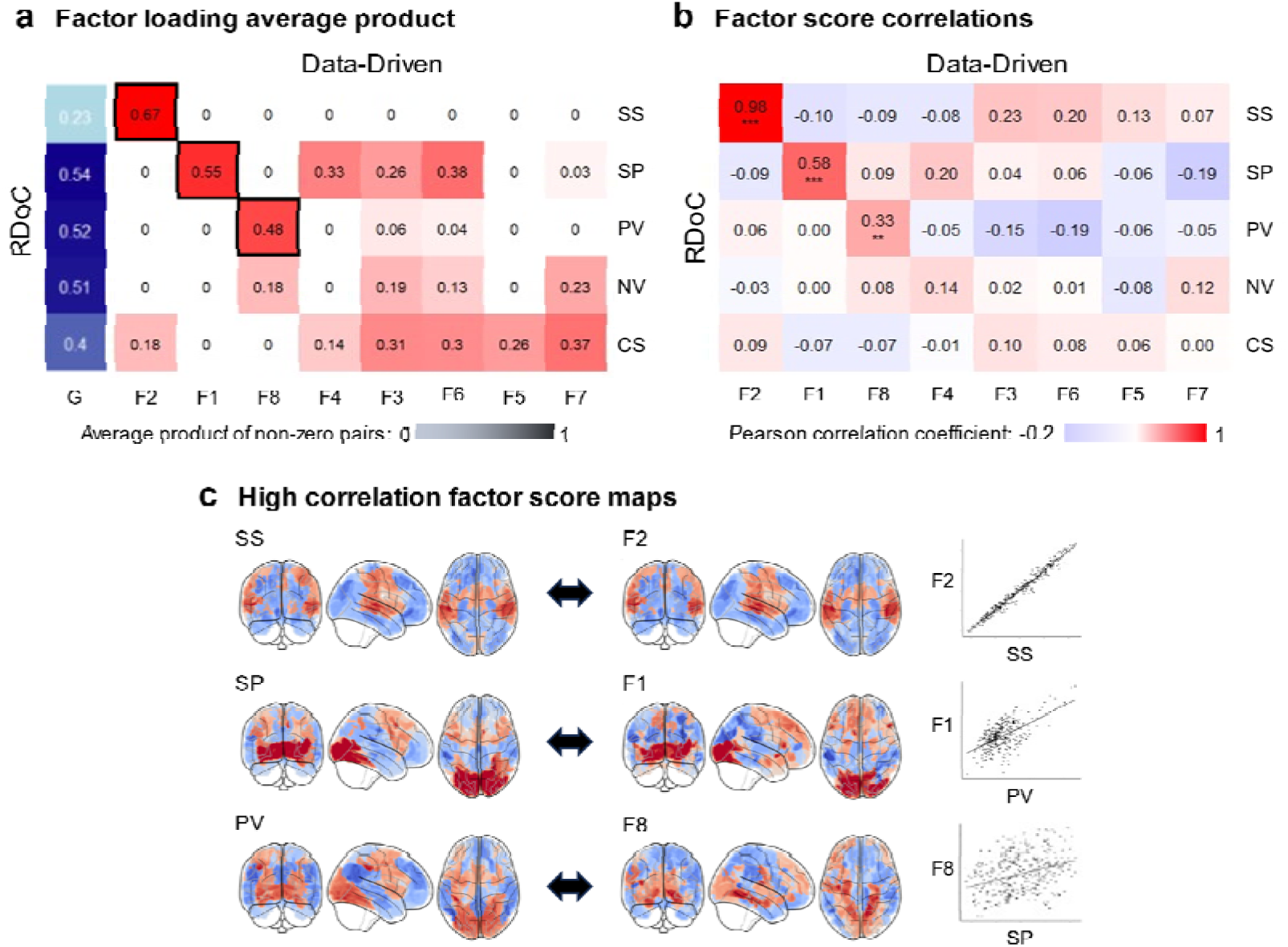
Factor convergence. (a) Heatmap showing the average product of factor loadings for maps in both the RDoC specific factor model and data-driven bifactor model. High values^20^ above ±.4 are highlighted with black borders. Maps of domains cross-loading on many specific factors reflect low intra-domain consistency; maps of domains sharing high loading with other domains on the same specific factor reflect high inter-domain similarity. (b) The heatmap shows the Pearson correlation of factor scores for the data-driven bifactor and RDoC specific factor model with asterisks representing p-values adjusted for spatial autocorrelation (***^/^\*\**p_adj_* ≤ .001-.003). Rows and columns are organized to show the strongest correlations in the diagonal. (c) Glass brains of factors with the highest correlations are shown as illustrative examples of strong one-to-one convergence in factor scores on the brain (warmer colors = positive scores; cooler colors = negative scores). Specifically, the sensorimotor systems domain displayed a strong correspondence with data-driven factor 2, the positive valence systems domain aligned with data-driven factor 1, and the social processes domain strongly correlated with data-driven factor 8. Scatter plots of correlations are shown on the right. CS: Cognitive Systems; NV: Negative Valence systems; PV: Positive Valence systems; SS: Sensorimotor Systems; SP: Social Processes; F1-6: specific factors 1-8.

The cognitive systems and negative valence systems domains load across multiple specific factors, indicating low intra-domain consistency. This suggests a degree of heterogeneity within the boundaries of these domains. In contrast, the sensorimotor systems domain shows notable intra-domain consistency by loading heavily on only a single data-driven factor (Fig. 4a), indicating a relatively consistent pattern in the activation maps of this domain. The positive valence systems and social processes domains demonstrate loadings across various data-driven factors, with particularly high loadings for data-driven factors 8 and 1, respectively. This implies that the boundaries of these domains may benefit from some refinement, given the observed complexities in their activation patterns across different factors. RDoC specific factors that share high loadings with data-driven factors (Fig. 4a) also show high factor score correlations (Fig. 4b-c)

Brain maps of factor scores and map loading for the data-driven bifactor and RDoC specific factor model are shown in Fig. 5. All of the RDoC domains but the sensorimotor systems domain show positive factor scores across both visual and motor regions, implicating the frequent recruitment of these regions across tasks of different domains. The sensorimotor systems domain predictably showed notable positive factor scores across the motor cortex. Similarly, the factors score brain map of the data-driven bifactor model’s general factor captured the predominant recruitment of visual and motor regions across most tasks. In contrast, the factor scores of the data-driven model’s specific factors captured more specific and varied functional activation patterns.

**Figure 5:**
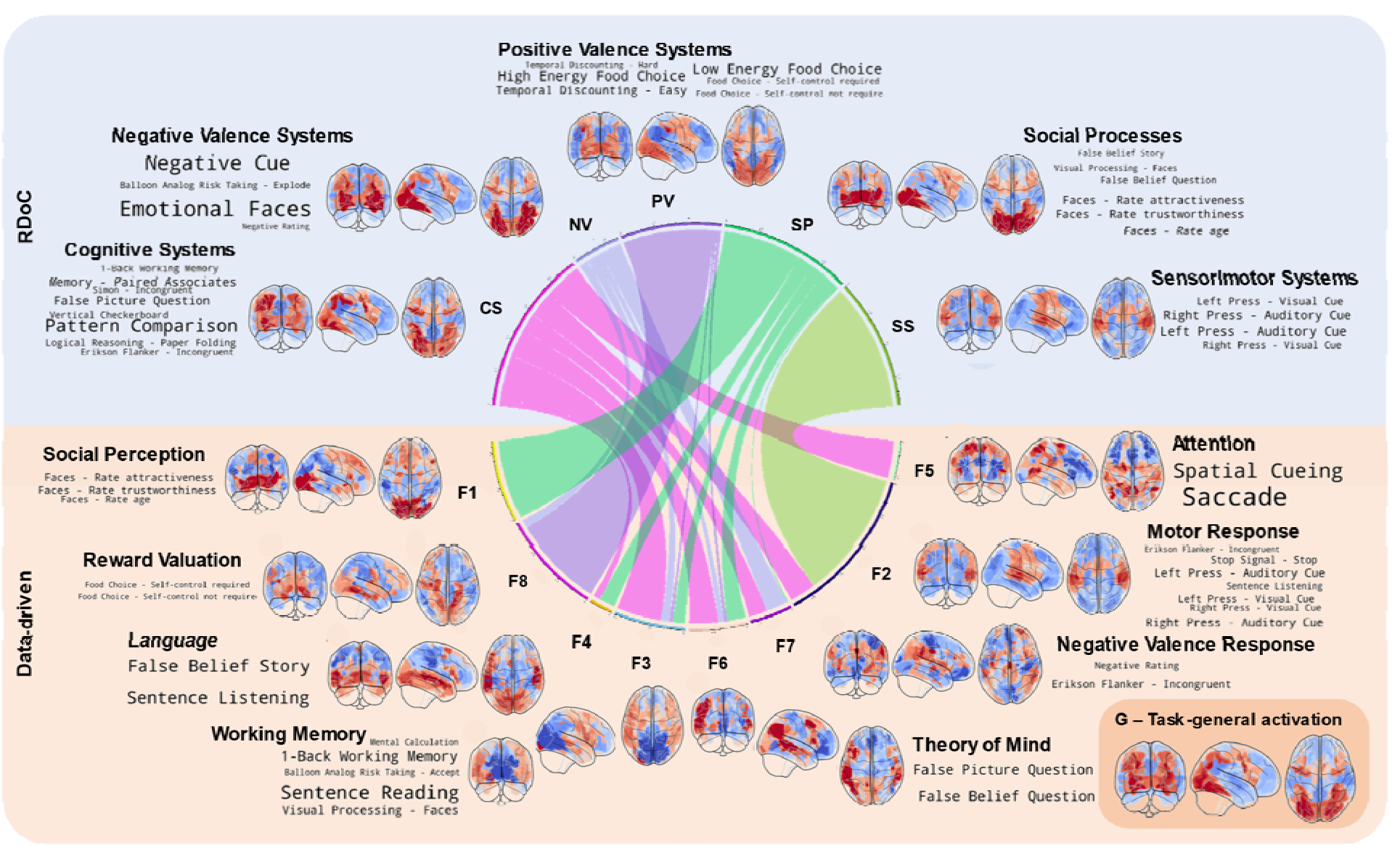
Mapping factors of the RDoC specific factor model and the data-driven bifactor model using data from whole-brain activation maps. Chord diagram showing the weighted links between maps loading on both models. The product of loadings in both models weighs links. Overlaps in maps showing loadings on both models illustrate the complex relations between RDoC factors and those derived using a data-driven bifactors modeling approach. Glass brain maps reflect factor scores (warmer colors = positive scores; cooler colors = negative scores). Word clouds of factors reflect the paradigm descriptors of the top eight maps loading on each factor. The size of words reflects the magnitude of the factor’s loading.

### Validation with held-out whole-brain activation maps and Neurosynth coordinate activation maps

We used a multi-prong validation strategy to assess the validity of the model solution derived from the curated training dataset. We compared the factor solutions from the RDoC specific factor model, representing the current RDoC framework, and the data-driven bifactor model, representing the best-performing data-driven model. For internal validation, we used the held-out maps from the original dataset, ensuring the model’s reliability within the same type of dataset (Fig. 6a). Additionally, we used Neurosynth coordinate activation maps that were a different data type (compared to whole-brain) and had better coverage of the RDoC domains (than the held-out maps) for external validation (Fig. 6b). This comprehensive validation strategy enabled us to evaluate the performance and generalizability of the factor structure we derived in varied contexts.

**Figure 6:**
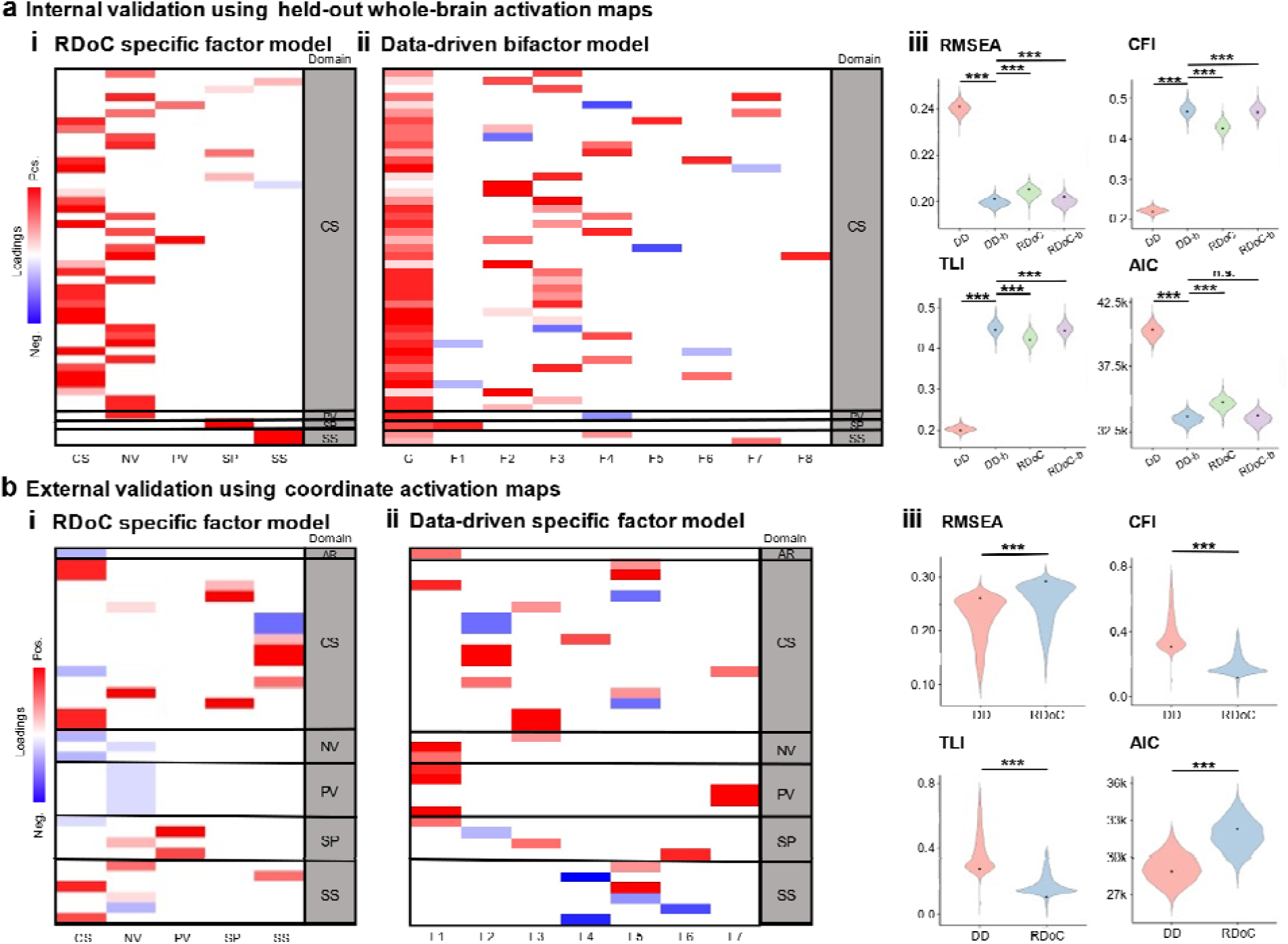
Validating RDoC and Data-driven models using held-out whole-brain and coordinate activation maps. Heatmaps showing factor loadings of the (i) RDoC and (ii) data-driven (DD) models for (a) whole-brain and (b) coordinate activation maps. Whole-brain activation maps held-out from the curated training dataset used to create the original models formed the internal validation set. The external validation set consists of coordinate activation maps obtained using Neurosynth’s LDA topic-based meta-analysis. These coordinate activation maps represent RDoC construct seed terms. The data-driven model for external validation with coordinate activation maps did not incorporate a general factor. This omission stemmed from the nature of sparse coordinate activation maps, which lacked significant overlaps that would warrant the representation of a general factor, as detailed in Supplementary Figure 2. Warmer colors = positive; Cooler colors = negative factor loadings shown. The held-out whole-brain and coordinate activation maps were assigned to specific factors based on factor scores from the data-driven and RDoC models from the training set. (iii) Relative fit measures for the different factor models. When applied to the held-out whole-brain activation maps, the data-driven bifactor model achieved the best overall fit, outperforming other models in most fit indices. The only exception was AIC, where the data-driven and RDoC bifactor models were tied as the best-fitting models. When applied to the coordinate activation maps, the data-driven specific factor model outperformed the RDoC specific factor model in fit scores. The violin plots represent the distribution of the bootstrap samples; the fit values from our models are indicated with black dots. AR: Arousal and Regulatory Systems; CS: Cognitive Systems; NV: Negative Valence systems; PV: Positive Valence systems; SS: Sensorimotor Systems; SP: Social Processes; G: General factor; F1-8: specific factors 1-8; DD: Data-driven specific factor model; DD-b: Data-driven bifactor model; RDoC: RDoC specific factor model; RDoC-b: RDoC bifactor model; n.s.: not significant.

For internal validation using held-out whole-brain activation maps, comparing the model fit of factors derived from the RDoC and data-driven models, our analysis revealed that the data-driven bifactor model exhibited the best fit for the held-out maps (robust RMSEA: *F*(3, 19996) = 425,763, *p* < .001; robust CFI: *F*(3, 19996) = 496,125, *p* < .001; robust TLI: *F*(3, 19996) = 467,909, *p* < .001; BIC: *F*(3, 19996) = 283,472, *p* < .001). The only exception was AIC (*F*(3, 19996) = 287,574, *p* < .001), where the data-driven and RDoC bifactor models were tied for best fit compared to the specific factor models. Unexpectedly, the data-driven specific factor model underperformed. Despite conducting additional model checks, no apparent errors in model fitting were identified. Tukey pairwise comparisons are shown in Supplementary Table 2.

External validation with Neurosynth coordinate activation maps was conducted to evaluate the model’s generalizability to diverse data types. We did not include a general factor in our data-driven model. Here, coordinate activation maps are sparse and do not show substantial overlaps that a general factor would represent. Indeed, the general factor of a data-driven bifactor model from a CFA exhibits limited loading across all the coordinate activation maps, indicating a lack of substantial influence (Supplementary Figure 2). The data-driven specific factor model demonstrated a better fit for the Neurosynth coordinate activation maps compared to the RDoC model (robust RMSEA: *t*(4171.3) = 16.1, *p* < .001; robust CFI: *t*(4116.6) = -41.9, *p* < .001; robust TLI: *t*(4094.0) = -38.9, *p* < .001; AIC: *t*(4168.3) = 70.3, *p* < .001; BIC: *t*(4168.3) = 69.1, *p* < .001). Taken together, these results indicate that the data-driven bifactor and specific factor models had generally better fit when generalized to unseen whole-brain and coordinate activation maps respectively compared to the RDoC models.

## Discussion

The current study aimed to advance the ontology of human brain functions by using a latent variable approach with bifactor analysis to examine the hierarchical structure of the RDoC framework. While it may be expected that data-driven models outperform a priori defined models, our study makes a unique contribution by validating this improved fit using previously unseen data and a different data modality (i.e., whole-brain vs. peak coordinate activation maps). This demonstrates the generalizability and robustness of our data-driven approach. Moreover, we provide concrete directions for refining the RDoC framework by delineating what the superior data-driven model could entail. These refinements are crucial for evolving the RDoC model in alignment with empirical data, ultimately enhancing the precision and applicability of psychiatric nosology.

The traditional RDoC model had most maps within each domain that loaded significantly onto each factor representing their domains; however, compared with data-driven models, the RDoC model also showed a relatively poor fit for both whole-brain and coordinate activation maps, indicating that the RDoC framework may not fully capture the complexity of brain-behavior relations. Adding a general factor to the conventional RDoC also improved the fit of the RDoC specific factor model, suggesting that the conventional RDoC framework may benefit by adding a superordinate domain representing task-general functioning. Incorporating a task-general functional domain into the RDoC model that extends beyond the existing task-specific functional domains would enhance the model’s ability to represent brain functioning comprehensively. However, the general factor capturing extensive visual cortex activation is likely influenced by the prevalence of visual stimuli in most fMRI tasks. Therefore, the general factor likely reflects a combination of the visual component of common fMRI tasks and a task-general functional domain.

Compared to the RDoC model, the data-driven model had a better fit to the data, indicating that it may provide a more accurate representation of the organization of circuit-function relations in the human brain. By differentiating general activation patterns common across different functional tasks from patterns specific to each construct, the data-driven bifactor model captured both shared and unique variance among different constructs, providing insight into the hierarchical organization of circuit-function relations. This is consistent with findings from recent studies that have advocated for a data-driven bifactor approach to understanding brain-behavior relations^17^. Notably, the data-driven specific factor model had better fitness scores after penalizing for model complexity as measured by both AIC and BIC. This indicates that although the data-driven bifactor model had the best overall model fit, the improvement in fit from adding the general factor comes at a substantial cost in model complexity.

The product matrix (Fig. 4a) and factor score correlations (Fig. 4b) revealed divergent patterns in correspondence across data-driven factors for different RDoC domains. For instance, whereas the cognitive systems domain had low loadings and correlations spread across the data-driven factors, the maps labeled by the positive valence systems, social processes, and sensorimotor systems domains had significant loadings and correlations confined to relatively fewer specific factors. Finally, the negative valence systems domain did not have significant loadings on any data-driven factors (Fig. 3). Still, its factor scores correlated strongly with two data-driven factors (Fig. 4). This pattern suggests that activation patterns within some domains are more distinct and separable than others, supporting our hypothesis that the boundaries between RDoC domains need to be reconsidered. Specifically, constructs within the cognitive systems domain might be better defined by being divided into separate domains. For example, attention, working memory, semantic processing/perception, and theory of mind within the cognitive systems domain formed individual data-driven factors (Fig. 5), and may be better represented as a revised set of domains in a refined RDoC framework.

Visualization of the factor scores on the brain showed us that factors from the RDoC models, excluding the sensorimotor system’s factor, consistently reveal activation patterns spanning visual and motor regions. This alignment with the general factor of the data-driven bifactor model suggests that there is shared task-general activation across tfMRI whole-brain activation maps. The utility of the general factor in the data-driven model lies in its ability to capture overarching patterns present across the entire dataset. This, in turn, allows the specific factors to focus on representing activation patterns that exhibit greater sensitivity to the nuances of specific task paradigms.

After constructing our factor models, we performed validation steps to assess how well our derived model, developed from the curated training dataset, could extend to unseen data. We used two distinct validation sets: whole-brain activation maps held out from the original dataset (internal validation) and coordinate activation maps sourced externally from Neurosynth (external validation). The internal validation using held-out whole-brain activation maps, while sharing the same data type as the original dataset, had a skewed distribution of maps (more cognitive maps) across the RDoC domains. To address this imbalance, we also conducted validation using Neurosynth coordinate activation maps, which provide a more balanced representation of the RDoC domains and constructs. This dual validation approach enhances the reliability of our findings and strengthens our model’s applicability to diverse datasets and contexts. Our data-driven bifactor model exhibited the best overall fit when applied to held-out whole-brain activation maps, though it tied with the RDoC bifactor model on AIC. For the coordinate activation maps, we excluded the general factor due to data sparsity, and the data-driven specific factor model showed superior fit for all measures compared to the RDoC model. These outcomes underscored the different data-driven model’s capability to capture brain activation patterns, extending beyond the initial dataset. Moreover, external validation using coordinate activation maps highlighted the data-driven model’s adaptability to diverse data types, particularly in handling sparse coordinate activation maps commonly generated from large meta-analytic tools.

Despite these advancements, it must be acknowledged that the overall model fit, even with the data-driven approaches, was not optimal. This limitation underscores the need for continued development in this field, recognizing that the complexity of brain-behavior relations may pose challenges to modeling efforts. While we curated a specific sample of whole-brain activation maps, this dataset represents a limited sample, with an imbalance in the representation of certain RDoC domains and constructs, and variability in the number of subjects across studies. This imbalance reflects the current focus areas and available datasets within the neuroimaging community. Future work should include a broader and more balanced range of tasks to increase the generalizability of our findings and examine the organization of the RDoC framework’s constructs. It is also important to note that although task activation relative to baseline allowed us to capture general task activation in our models, tasks often involve more than one functional domain. For example, even a simple button press-to-cue task involves perception (cognitive domain) and motor action (sensorimotor systems domain). Therefore, subtraction contrasts between tasks and other model structures may reveal additional insights into the brain’s function-circuit relations.

While the RDoC framework was designed top-down as a conceptual framework to integrate multiple levels of analysis, its different levels of analysis have not been substantially validated empirically. Our study uses tfMRI data to provide insights into the framework’s performance, specifically within the context of brain circuit-function relations. Future research should continue to evaluate the RDoC framework across other units of analysis, such as genetic and behavioral data, to inform further refinements and to ensure its comprehensive applicability.

In conclusion, our study indicates that a data-driven approach provides a more accurate representation of the organization of the human brain’s circuit-function relations than the conventional RDoC model. Our findings support the use of data-driven approaches to inform revisions to the RDoC framework and to develop a more comprehensive ontology to guide further research. Integrating a task-general domain within the RDoC framework holds promise in broadening the capacity of the RDoC framework to capture brain functionality holistically. Furthermore, our research underscores the need to reassess the demarcations or boundaries within RDoC domains. However, this is not the only path forward. An integrative approach may be necessary, including developing new neuroimaging techniques and tasks to address currently underrepresented domains. The end goal is not solely to adopt a data-driven model or refine RDoC, but rather, to advance conceptual and empirical methods. We acknowledge that an effective solution is likely to be more complex than simply amending the RDoC framework and will require a multifaceted strategy.

## Methods

### Gathering and preparing activation maps

Our dataset comprises both whole-brain activation maps and maps reconstructed from activation coordinates (Fig. 1). These two sets of maps capture the primary published forms of neuroimaging data. Whole-brain activation maps underwent a rigorous selection process due to variations in contrast methodologies and acquisition parameters. Only maps from healthy participants were included. The following subsections describe how we gathered and processed these maps.

#### Whole-brain activation maps

The collection of 84 whole-brain tfMRI maps was curated by Bolt et al.^17^ and sourced from two publicly accessible datasets: Neurovault^21^(n=82) and UK Biobank^22^ (n=2). Although maps were also sourced from the Human Connectome Project by Bolt et al.^17^, these maps did not correlate sufficiently strongly with the other 84 maps within the dataset and, consequently, were not included for further analysis (Supplementary Figure 3). We used only unthresholded group-level BOLD contrasts for task conditions versus baseline. Contrast maps corresponding to the subtraction between two activation maps were not included because contrasts between events within the task would eliminate general activation patterns representing the task’s domain.

We categorized contrast maps by matching the task descriptions extracted from the associated task contrasts (e.g., from https://neurovault.org/ for NeuroVault) with descriptions of the RDoC domains and construct definitions from the RDoC matrix^23^ (Supplementary Table 1). For instance, a contrast map created from a task where participants press a button as directed by visual instructions is categorized under the sensorimotor domain. We restricted our analysis to the following RDoC domains: cognitive systems, positive valence systems, negative valence systems, social processes, and sensorimotor systems, as no activation maps in the dataset fit within the domain of arousal and regulatory systems. This gap underscores the need for more future research to design and include tasks that specifically target the arousal and regulatory systems domain. Recognizing that a substantial proportion of the activation maps in the initial dataset originated from the cognitive systems domain (70%), we curated a sub-collection of maps. The curated training dataset was designed to achieve a more balanced representation of the constructs across all five domains and minimize study overlap. The curated training dataset is composed of 37 maps derived from a total of 6,119 participants, distributed as follows: cognitive systems (40.5%), negative valence systems (13.5%), positive valence systems (18.9%), social processes (16.2%), and sensorimotor systems (10.8%). We also excluded maps representing tasks that strongly implicated multiple RDoC domains. Details and sex balance of all 37 curated training maps and the 47 held-out maps (used for the internal validation set) are listed in (Supplementary Table 1).

Our initial collection of maps was composed of both *t*-stat and *z*-stat images. The unthresholded *t*-stat images were first converted to *z*-stat images before further processing. All maps were then resampled to the 2mm MNI-152 standard-space T1-weighted template (Nonlinear 6th generation).

#### Map post-processing

All activation maps were parcellated into 333 cortical and 14 subcortical brain regions using the Gordon^24^ and Harvard-Oxford^25^ atlases, respectively. Before performing factor analysis, the activation values for each map were also scaled to minimize the effects of varying acquisition parameters across different studies and to enhance the convergence of the factor models.

### Factor analysis

Data-driven models encompassed an exploratory factor analysis (EFA) step to first identify potential factor structures, followed by a confirmatory factor analysis (CFA) step to assess how well the factor model fits the observed data. In contrast, RDoC models involved only a CFA step, given that they incorporated pre-defined factors specific to RDoC.

Bootstrap distributions of fit indices were computed by resampling parcels over 5,000 iterations. Factor scores were estimated using Bartlett’s method to create brain maps (Fig. 5) reflecting each region’s loading for each factor. This method is designed to yield factor scores that are strongly correlated with their respective factor, while maintaining minimal or no correlation with other factors.

#### Data-driven Factor Analysis with whole-brain activation maps

The factor analysis for our data-driven models (Fig. 2, bi and bii) was composed of three primary stages: (1) Horn’s parallel analysis to determine the optimal number of factors to extract (see below); (2) EFA to extract specific factors; and (3) CFA with both the specific factors and a general factor for the bifactor model, and only specific factors for the specific factor model.

To determine the number of factors to extract, we conducted parallel analysis^26^, which identifies the number of factors to extract based on where the calculated eigenvalues of the actual data intersect with the eigenvalues of random data generated^27^. We then conducted an EFA using principal axis factoring and oblimin rotation to extract the identified number of specific factors in the subsequent confirmatory analysis. We also examined the scree plots to verify the suitability of the number of factors extracted (Supplementary Figure 1). To conduct the EFA, we used oblimin rotation to allow for correlated factors, but the correlation was constrained to be small. Based on previous work, each specific factor was defined by maps with a high absolute loading of 0.4 or higher^20^. For the CFA, we used robust maximum likelihood estimation to account for non-normality in the data. Orthogonal rotation was used in the bifactor models to ensure that the general factor is not contaminated by the specific factors, making it difficult to interpret the factor structure. By constraining the general factor to be orthogonal to the specific factors, bifactor models can identify a general factor independent of the specific factors. The general factor captures the shared variance, while the specific factors capture the distinct variances that are unique to subsets of activation maps^28^. We used the specific factors from the EFA and a general factor with all maps loaded onto it. For comparison, we also conducted an alternate CFA without the general factor (specific factor model). To account for interrelationships between factors within the specific factor models, which are not captured by a general factor, we maintained non-orthogonality and allowed all of our specific factor models to exhibit covariance.

#### RDoC Domain Factor Analysis with whole-brain activation maps

Our curated training set of whole-brain activation maps was grouped into RDoC domain-specific factors by matching the task description with the domain/construct definition. For our RDoC models (Fig. 2, ai and aii), we conducted a CFA utilizing robust maximum likelihood estimation and non-orthogonal factors.

### Validation using unseen data

To evaluate the robustness and generalizability of our model solutions, we conducted a validation procedure using both the held-out maps from the original dataset (internal validation) and the coordinate activation maps sourced from Neurosynth (external validation) (Fig. 6).

#### Internal Validation using held-out whole-brain activation maps

We systematically assigned individual maps to specific factors from the RDoC specific factor model (representing the RDoC framework) and the data-driven bifactor model. Factor assignment and loadings were determined using the factor scores derived from the original model using the curated training dataset. The factor assignment involved identifying, for each map, the factor from the original factor model that exhibited the highest product sum. After the map was assigned to a factor, the loading for each map was determined by dividing the map’s product sum for that factor by the highest product sum of all other maps that were assigned to the same factor, providing an adjusted coefficient for its association with the respective factor (details of the factor assignment process is shown in Supplementary Figure 4). Subsequently, we conducted a CFA with these factor assignments and loadings. We then compared the fit scores obtained from this validation analysis. This process allowed us to evaluate how well the training model solution generalized to unseen data, effectively probing the model’s capability to capture brain activation patterns beyond the curated training dataset.

#### External validation using Neurosynth coordinate activation maps

In addition to using the held-out maps from our initial dataset to test the model solution derived using the curated dataset, we also utilized coordinate activation maps with topics matching RDoC construct seed terms for external validation. Seed terms adapted from Beam et. al.^7^ were compiled based on the name and synonyms of each RDoC domain construct, e.g., “acute threat” and “fear” for the “acute threat” construct under the negative valence system domain. These seed terms were then used to search for matching terms in a topic-based meta-analysis using Neurosynth. 200 topics were extracted using Latent Dirichlet Allocation (LDA) from the abstracts of all articles in the latest version of Neurosynth^9^ (ver. 5). Neurosynth’s LDA topic-based meta-analysis is a data-driven approach that uses natural language processing (NLP) techniques to uncover topics that share terms across a large set of studies. Each topic is associated with a probabilistic reverse inference map representing the likelihood that a given brain coordinate is activated during a study using these terms. Using this meta-analysis technique, we identified 36 coordinate activation maps with topics that matched RDoC construct seed terms. Seed terms with multiple topic maps had their activation averaged before further analysis. Spatial smoothing was applied using a 12-mm full-width half-maximum (FWHM) Gaussian kernel centered on each peak-activation coordinate in the maps, creating more realistic representations of brain activation patterns. Values were then thresholded (*z* > 0.1) to remove noise from using a Gaussian kernel. A complete list of the seed terms and topics sourced from Neurosynth is presented in Supplementary Table 3. These maps were then used to validate the factor structure from the curated training dataset in the same way as the held-out whole-brain activation maps.

### Statistical Analysis

To assess the potential influence of sex balance, sample size, and other differences across studies, such as acquisition parameters, on our model, we conducted regression analyses with these variables using the curated dataset. The sex balance, represented as the ratio of males to females, showed an average explained variance (*R*^2^) of 5.8% (*SD* = 8.2%) across brain regions. Only activation in 6.34% of brain regions had a significant linear relation with sex balance (*p_fdr_* < .05) after adjusting for multiple comparisons. Even though only a small percentage of brain regions had a significant relation with sex balance, regressing out the effect of sex balance from the dataset caused substantial issues in the EFA stage. The correlation matrix was non-positive definite, meaning it could not be inverted as required for factor analysis, necessitating smoothing and leading to approximations in factor score estimates. In addition, the CFA failed to converge.

We also investigated the effect of sample size as a covariate that may affect the dataset. Sample size had an average *R*^2^ of 1.4% (*SD* = 2.2%) across brain regions, with no significant linear relation observed between activation and sample size in any brain regions (*p_fdr_* < .05) after adjusting for multiple comparisons. Similarly, regression analysis with study ID as a categorical variable, representing general differences in study parameters, indicated an average *R*² of 3.9% (*SD* = 4.6%) across brain regions, with no significant linear relationship observed (*p_fdr_*< .05). Based on these findings and consistent with Bolt et al’s^17^ initial approach with this dataset, we did not include sex, sample size, or study ID as covariates in our analysis.

Model fit was assessed using robust variants of fit indices, including the Root Mean Square Error of Approximation (RMSEA), Comparative Fit Index (CFI), and Tucker-Lewis Index (TLI). The robust variant of these fit indices were chosen to account for potential non-normality in the data. Additionally, information theoretical measures of model complexity, the Akaike Information Criterion (AIC) and Bayesian Information Criterion (BIC), were used for comparison. AIC and BIC consider the trade-off between model fit and complexity, with lower values indicating a more optimal balance^29^. The bootstrap distribution of these fit indices were computed using the Yuan bootstrap method^30^ of resampling with 5,000 iterations. With the Yuan bootstrapping method, the data is transformed by combining data and the model, such that the resampling space is closer to the population space. For the comparison of multiple models’ fit scores, Analysis of Variance (ANOVA) tests were employed. Subsequent post hoc pairwise comparisons were performed using Tukey’s Honestly Significant Difference test to determine the model with the best fit score. When comparing fit scores between only two models, t-tests were utilized instead.

Pearson correlations were calculated between the factor scores of the RDoC specific factor model and the data-driven bifactor model. This analysis aimed to explore the extent to which the loadings of the base RDoC model align with the factors derived from the data-driven approach. To account for spatial autocorrelation in our analyses, we utilized spatial autocorrelation-preserving surrogate maps generated with BrainSMASH (Brain Surrogate Maps with Autocorrelated Spatial Heterogeneity) as described by Burt et al.^31^ to calculate adjusted p-values (Supplementary Table 4). For this calculation, distances between brain regions were computed using Euclidean distances of MNI centroids for each parcel. All statistical tests were two-tailed.

## Data availability

The whole-brain task fMRI contrast maps used in this study are publicly available at the neurovault.org website. The coordinate activation maps used are available at neurosynth.org.

## Code availability

The R code used for latent variable analysis and visualization is available at https://github.com/braindynamicslab/rdoc-lfa

## Acknowledgments

This work was supported by an NIH R01MH127608 and an MCHRI Faculty Scholar Award to M.S.

## Author contributions

S.K.L.Q.: conceptualization, methodology, software, validation, investigation, visualization, and writing.

B.J.: methodology, validation, writing, and supervision.

C.G.: data curation for the peak coordinate activation maps from Neurosynth, visualization, and writing.

L.Q.U.: interpretation, writing, and supervision.

J.A.M.: interpretation, writing, and supervision.

D.M.B.: interpretation, writing, and supervision.

D.A.F.: interpretation, writing, and supervision.

I.H.G.: interpretation, writing, and supervision.

R.A.P.: interpretation, writing, and supervision.

M.S.: conceptualization, methodology, software, validation, interpretation, writing, and supervision.

