## Supplementary Information for "A Data-Driven Latent Variable Approach to Validating the Research Domain Criteria Framework"

Supplementary Figure 1. The scree plot resulting from the parallel analysis used in the data-driven exploratory factor analysis reveals that models with eight or fewer factors exhibit eigenvalues exceeding those anticipated by chance.

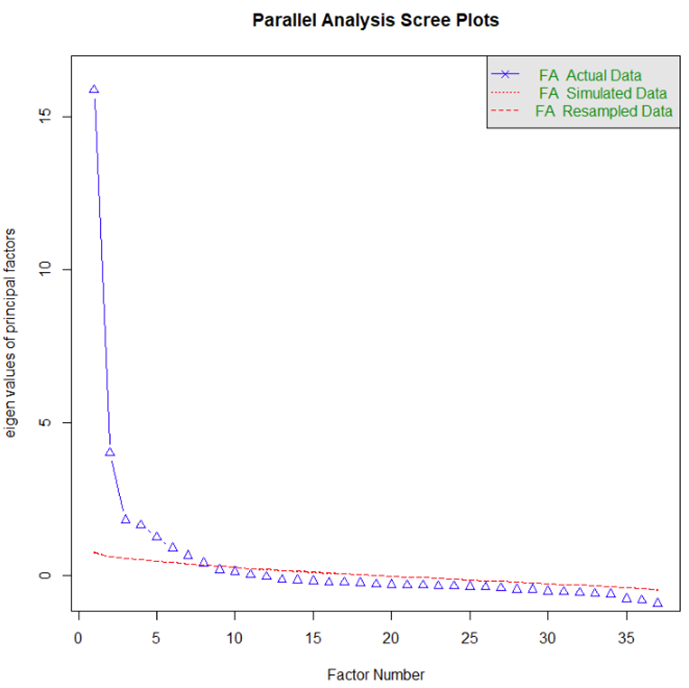

Supplementary Figure 2. The general factor, G of a data-driven bifactor model from a CFA exhibits limited loading across the coordinate maps, indicating a lack of substantial influence. G: General factor; F1-8: specific factors 1-8.

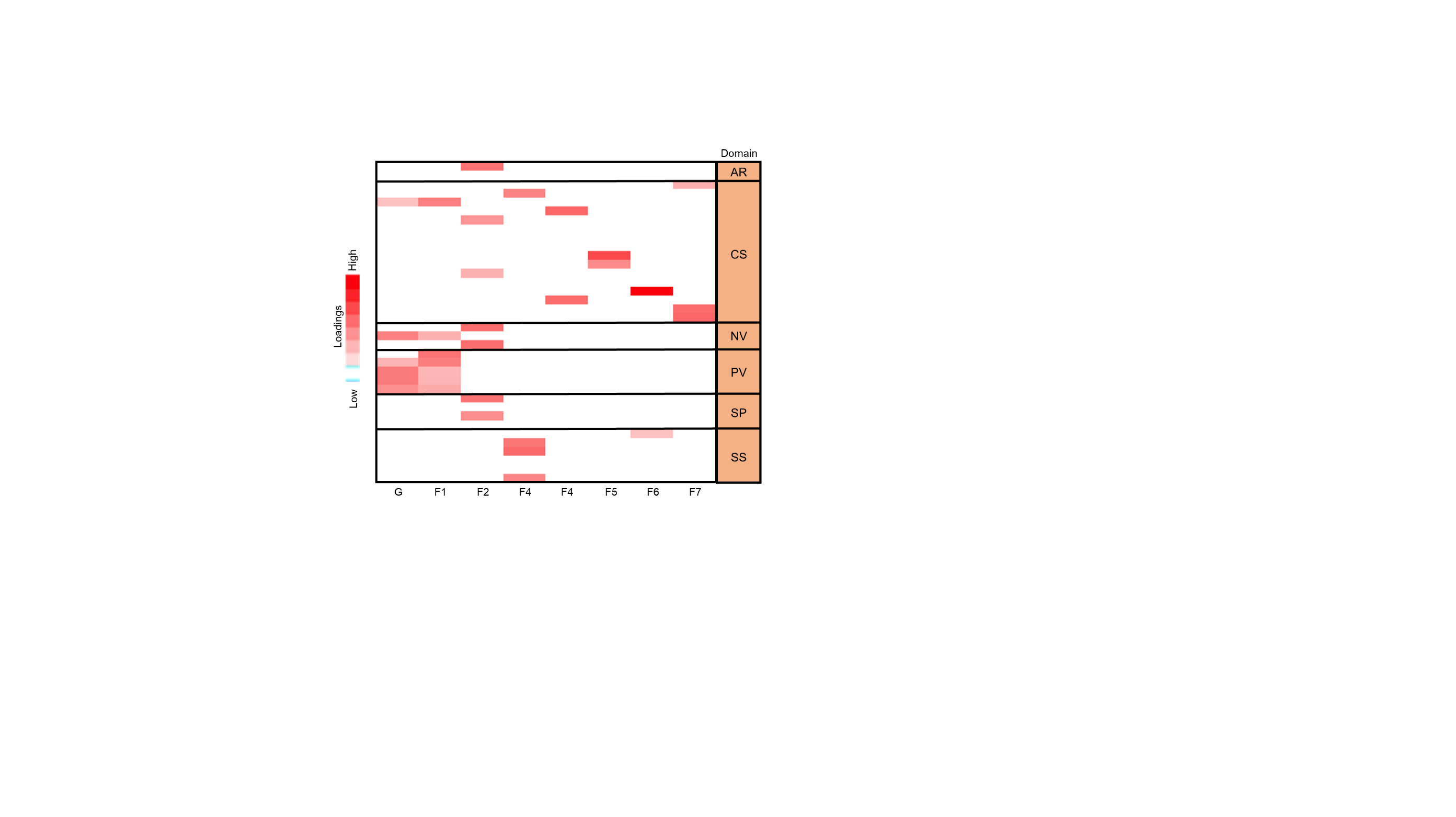

Supplementary Figure 3: The whole-brain activation maps derived from the Human Connectome Project (HCP) show no significant correlation with whole-brain activation maps outside of the HCP dataset. The following heatmap shows the Pearson correlation coefficients, *r* between all whole-brain activation maps. The low level of covariance between non-HCP and HCP maps suggests that HCP maps may not be suitable for integration with non-HCP maps in factor analysis.

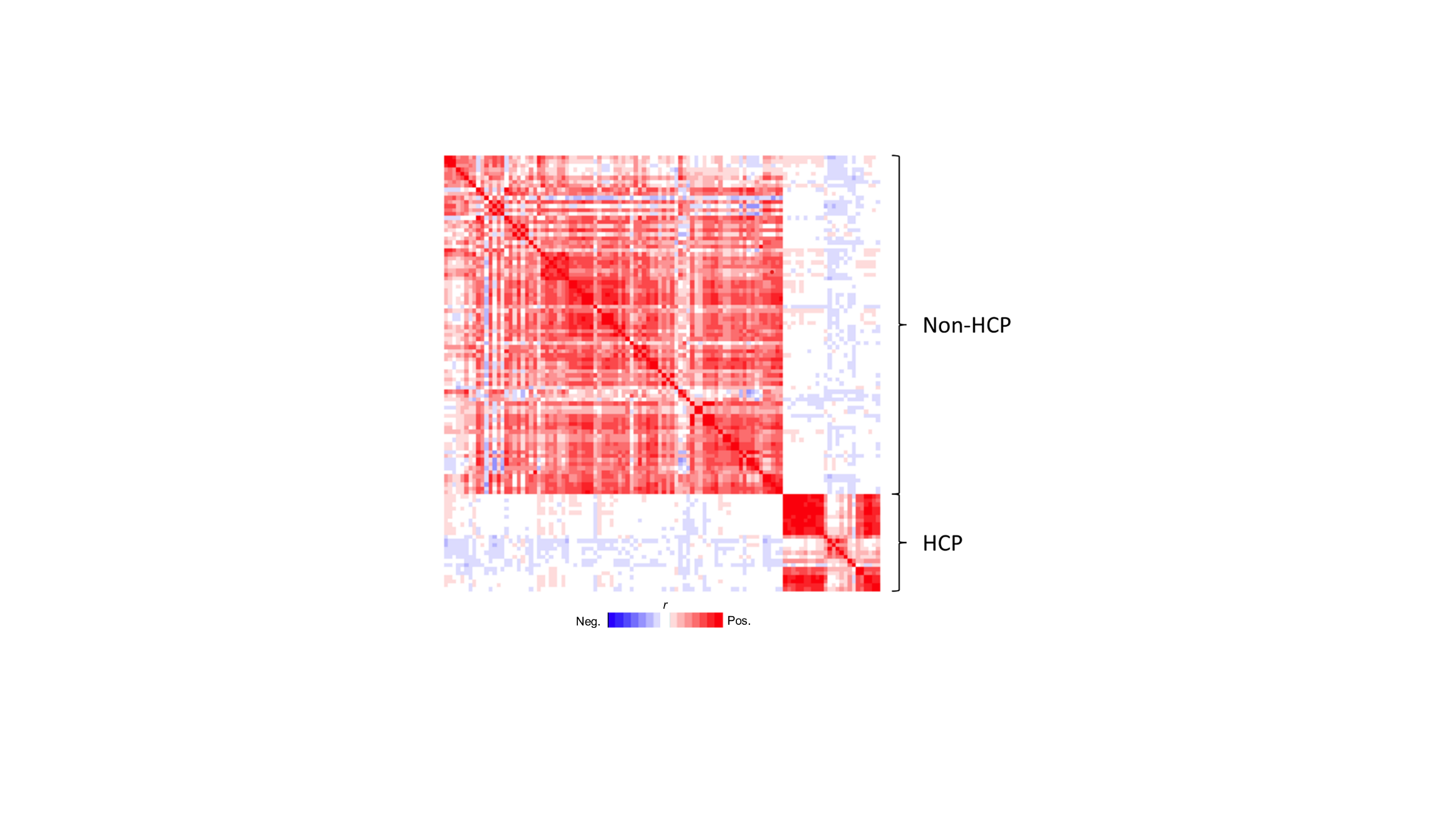

Supplementary Figure 4: Procedure for the derivation of the validation sets’ CFA models using specific factor scores from the training set and maps from the validation sets.

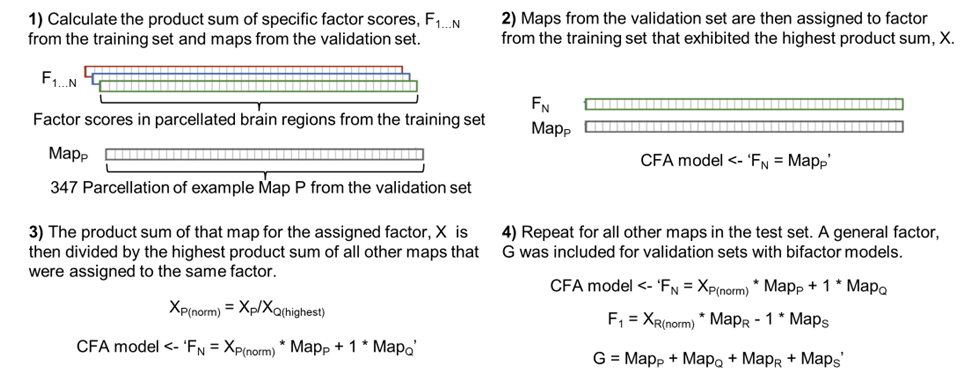

Supplementary Table 1: Table of all whole-brain maps in the (A) curated training and (B) test dataset. Details about the task were extracted from the associated websites (e.g., https://neurovault.org/ for Neurovault). (C) Table for the sex ratio for each study. N: Neurovault; B: Biobank; CS: Cognitive Systems; NV: Negative Valence systems; PV: Positive Valence systems; SS: Sensorimotor Systems; SP: Social Processes.

(A)

| No. | Source | ID | *N* | Domain | Task Paradigm | Condition | Details |
| --- | --- | --- | --- | --- | --- | --- | --- |
| 1 | N | 1212 | 23 | CS | Spatial cueing | Mean of Congruent + Incongruent | Look at the target after fixation if you get a cue |
| 2 | N | 45 | 26 | CS | Erikson Flanker | Incongruent, Correct | Press the target button while distractors flank it |
| 3 | N | 657 | 32 | CS | Eye movement | Saccade | Saccade to target |
| 4 | N | 98 | 24 | CS | Stop Signal | Successful Stop | Inhibit response |
| 5 | N | 99 | 21 | CS | Simon | Incongruent, Correct | The box appears on a different side as button press |
| 6 | N | 426 | 34 | CS | False Belief | False Picture Question | Question about outdated representations (control) |
| 7 | N | 857 | 303 | CS | Semantic Memory | Paired Associates | Remember paired words |
| 8 | N | 654 | 40 | CS | Sentence Reading | Normal Sentences | Read sentences |
| 9 | N | 94 | 94 | CS | Functional Localizer | Sentence Listening | Listen to a sentence |
| 10 | N | 1126 | 44 | CS | Lexical Decision | Fixation | Control condition, passive fixation |
| 11 | N | 857 | 303 | CS | Pattern Comparison | Pattern | Determine if the pattern matches |
| 12 | N | 94 | 94 | CS | Functional Localizer | Vertical Checkerboard | View flashing checkerboard |
| 13 | N | 46 | 49 | CS | 1-Back Working Memory | Objects | Determine if the target matches 1 stimulus before |
| 14 | N | 657 | 32 | CS | Mental Calculation | Calculation | Subtract using the previously shown number |
| 15 | N | 857 | 303 | CS | Logical Reasoning | Paper Folding | Select a pattern that would result from folding |
| 16 | N | 425 | 33 | NV | Emotion regulation | Negative Cue | View negative valenced image |
| 17 | N | 425 | 33 | NV | Emotion regulation | Negative Rating | Rate the negative valenced image |
| 18 | N | 98 | 24 | NV | Emotion regulation | Negative Cue | View negative valenced image |
| 19 | B | 2 | 5285 | NV | Face emotion | Emotional Faces | Which angry or fearful face matches the top face |
| 20 | N | 98 | 24 | NV | Balloon Analog Risk-taking | Explode | Balloon explodes |
| 21 | N | 724 | 20 | PV | Food Choice | Self-control not Required | Choose between high and low energy snack |
| 22 | N | 723 | 20 | PV | Food Choice | High Energy Food Choice | Whether they want to eat a high-energy snack |
| 23 | N | 724 | 20 | PV | Food Choice | Self-control Required | Choose between high tasty and low-energy snack |
| 24 | N | 723 | 20 | PV | Food Choice | Low Energy Food Choice | Whether they want to eat a low-energy snack |
| 25 | N | 98 | 24 | PV | Temporal Discounting | Hard Trials | Hard to pick an immediate or delayed reward |
| 26 | N | 98 | 24 | PV | Balloon Analog Risk-taking | Accept | Pump balloon for greater reward at risk of exploding |
| 27 | N | 98 | 24 | PV | Temporal Discounting | Easy Trials | Easy to pick an immediate or delayed reward |
| 28 | N | 426 | 34 | SP | False Belief | False Belief Question | Question about the story of a person's false belief |
| 29 | N | 426 | 34 | SP | False Belief | False Belief Story | Read the story of a person's false belief |
| 30 | N | 110 | 44 | SP | Social Judgements of Faces | Rate Attractiveness | Determine which face is more attractive |
| 31 | N | 110 | 44 | SP | Social Judgements of Faces | Rate Age | Determine which face is older |
| 32 | N | 110 | 44 | SP | Social Judgements of Faces | Rate Trustworthiness | Determine which face is more trustworthy |
| 33 | N | 657 | 32 | SP | Visual Processing | Faces | Visual stimuli presented |
| 34 | N | 94 | 94 | SS | Functional Localizer | Left Press - Auditory | Follow auditory instruction |
| 35 | N | 94 | 94 | SS | Functional Localizer | Left Press - Visual | Follow visual instruction |
| 36 | N | 94 | 94 | SS | Functional Localizer | Right Press - Auditory | Follow auditory instruction |
| 37 | N | 94 | 94 | SS | Functional Localizer | Right Press - Visual | Follow visual instruction |

(B)

| No. | Source | ID | *N* | Domain | Task Paradigm | Condition | Details |
| --- | --- | --- | --- | --- | --- | --- | --- |
| 1 | N | 98 | 24 | PV | Balloon Analog Risk-taking | Reject | Pump balloon for greater reward at risk of explode |
| 2 | N | 45 | 26 | CS | Erikson Flanker | Congruent, Correct | Press target button (not flanked by distractors) |
| 3 | B | 2 | 5285 | CS | Emotional face | Shapes (control) | Match fearful face/shape with other face/shape |
| 4 | N | 423 | 20 | CS | Stop Signal | Manual Response | All manual responses |
| 5 | N | 423 | 20 | CS | Stop Signal | Spoken Letter Naming Response | Response in spoken letter naming condition |
| 6 | N | 423 | 20 | CS | Stop Signal | Spoken Pseudoword Naming Response | Response in spoken pseudoword naming condition |
| 7 | N | 98 | 24 | CS | Stop Signal | Go | Press directional button as shown |
| 8 | N | 425 | 33 | CS | Emotion regulation | Reappraise Negative Cue | Reappraise negative valenced image |
| 9 | N | 98 | 24 | CS | Emotion regulation | Suppress Negative Cue | Suppress negative valenced image |
| 10 | N | 99 | 21 | CS | Simon | Incongruent, Incorrect | Box appears on different side as button press |
| 11 | N | 857 | 303 | CS | Vocabulary | Antonyms | Identify the antonym |
| 12 | N | 857 | 303 | CS | Vocabulary | Picture Naming | Name the picture |
| 13 | N | 857 | 303 | CS | Vocabulary | Synonyms | Identify the synonym |
| 14 | N | 857 | 303 | CS | Semantic Memory | Logical Memory | Remember story details |
| 15 | N | 857 | 303 | CS | Semantic Memory | Word Order | Remember word order |
| 16 | N | 657 | 32 | CS | Auditory Processing | French Words | Auditory stimuli presented |
| 17 | N | 657 | 32 | CS | Auditory Processing | Korean Words | Auditory stimuli presented |
| 18 | N | 25 | 32 | CS | Semantic Priming | Letter Strings | Presented meaningless letter strings |
| 19 | N | 654 | 40 | CS | Sentence Reading | Pseudo Sentences | Read jabberwocky sentences |
| 20 | N | 657 | 32 | CS | Visual Processing | Words | Visual stimuli presented |
| 21 | N | 657 | 32 | CS | Auditory Processing | Sound | Auditory stimuli presented |
| 22 | N | 425 | 33 | CS | Emotion regulation | Neutral Cue - Visual | View neutral image |
| 23 | N | 98 | 24 | CS | Emotion regulation | Neutral Cue - Visual | View neutral image |
| 24 | N | 94 | 94 | CS | Functional Localizer | Horizontal Checkerboard | View flashing checkerboard |
| 25 | N | 857 | 303 | CS | Pattern Comparison | Digit symbol | Determine if pattern matches |
| 26 | N | 657 | 32 | CS | Visual Processing | Action | Action in response to stimuli |
| 27 | N | 657 | 32 | CS | Visual Processing | Digit | Visual stimuli presented |
| 28 | N | 657 | 32 | CS | Visual Processing | House | Visual stimuli presented |
| 29 | N | 657 | 32 | CS | Visual Processing | Scramble | Visual stimuli presented |
| 30 | N | 657 | 32 | CS | Visual Processing | Tool | Visual stimuli presented |
| 31 | N | 94 | 94 | CS | Functional Localizer | Calculation, Auditory | Perform mental calculation |
| 32 | N | 98 | 24 | CS | Emotion regulation | Rating | Rate negative/neutral image seen |
| 33 | N | 426 | 34 | CS | False Belief | False Picuture Story | Read story of outdated physical representations |
| 34 | N | 94 | 94 | CS | Functional Localizer | Calculation, Visual | Perform mental calculation |
| 35 | N | 857 | 303 | CS | Logical Reasoning | Letter Set | Select which letter groups is different from others |
| 36 | N | 857 | 303 | CS | Logical Reasoning | Matrix Reasoning | Select which pattern best completed matrix |
| 37 | N | 446 | 21 | CS | Match to sample | Matching Task | Which shape matched target |
| 38 | N | 857 | 303 | CS | Pattern Comparison | Letter | Determine if strings match |
| 39 | N | 46 | 49 | CS | 1-Back Working Memory | Consonant Strings | Determine if target matches 1 stimulus before |
| 40 | N | 46 | 49 | CS | 1-Back Working Memory | Scrambled Objects | Determine if target matches 1 stimulus before |
| 41 | N | 46 | 49 | CS | 1-Back Working Memory | Words | Determine if target matches 1 stimulus before |
| 42 | N | 94 | 94 | CS | Functional Localizer | Sentence Reading | Read sentence |
| 43 | N | 99 | 21 | CS | Simon | Congruent, Correct | Box appears on same side as button press |
| 44 | N | 99 | 21 | CS | Simon | Congruent, Incorrect | Box appears on same side as button press |
| 45 | N | 110 | 44 | SS | Social Judgements of Faces | Left Motor Response | Left face is answer to judgment |
| 46 | N | 110 | 44 | SS | Social Judgements of Faces | Right Motor Response | Right face is answer to judgment |
| 47 | N | 110 | 44 | SP | Social Judgements of Faces | Rate Happiness | Determine which face is happier |

(C)

| No. | Source | ID | % Sex (M/F) |
| --- | --- | --- | --- |
| 1 | N | 45 | 61.5/38.5 |
| 2 | N | 46 | 51.1/48.9 |
| 3 | N | 94 | 47.9/52.1 |
| 4 | N | 98 | 58.3/41.7 |
| 5 | N | 99 | 57.1/42.9 |
| 6 | N | 110 | 54.5/45.5 |
| 7 | N | 425 | 40.0/60.0 |
| 8 | N | 426 | 38.7/61.3 |
| 9 | N | 654 | 57.5/42.5 |
| 10 | N | 657 | 100/0 |
| 11 | N | 723 | 0/100 |
| 12 | N | 724 | 0/100 |
| 13 | N | 857 | 45.9/54.1 |
| 14 | N | 1126 | 29.5/70.5 |
| 15 | N | 1212 | 0/100 |
| 16 | B | - | 47.0/53.0 |

Supplementary Table 2. Tables of all Tukey test pairwise comparisons for fit indices from the data-driven and RDoC models for (A) the curated dataset and (B) the internal validation step.

**A.**

**Table for pairwise comparison of RMSEA Fit Measure**

| Comparison | Estimate | Std. Error | t-value | p-value |
| --- | --- | --- | --- | --- |
| Data-driven - Data-driven (bifactor) | 0.02522 | 5.58e-05 | 452.0 | <.001*** |
| RDoC - Data-driven (bifactor) | 0.02859 | 5.58e-05 | 512.4 | <.001*** |
| RDoC (bifactor) - Data-driven (bifactor) | 0.01254 | 5.58e-05 | 224.8 | <.001*** |
| RDoC - Data-driven | 0.00337 | 5.462e-05 | 61.7 | <.001*** |
| RDoC (bifactor) - Data-driven | -0.01268 | 5.462e-05 | -232.2 | <.001*** |
| RDoC (bifactor) - RDoC | -0.01605 | 5.462e-05 | -293.9 | <.001*** |

**Table for pairwise comparison of CFI Fit Measure**

| Comparison | Estimate | Std. Error | t-value | p-value |
| --- | --- | --- | --- | --- |
| Data-driven - Data-driven (bifactor) | -0.043815 | 2.341e-04 | -187.2 | <.001*** |
| RDoC - Data-driven (bifactor) | -0.139133 | 2.341e-04 | -594.4 | <.001*** |
| RDoC (bifactor) - Data-driven (bifactor) | -0.049895 | 2.341e-04 | -213.2 | <.001*** |
| RDoC - Data-driven | -0.095317 | 2.292e-04 | -416.0 | <.001*** |
| RDoC (bifactor) - Data-driven | -0.006080 | 2.291e-04 | -26.5 | <.001*** |
| RDoC (bifactor) - RDoC | 0.089237 | 2.291e-04 | 389.5 | <.001*** |

**Table for pairwise comparison of TLI Fit Measure**

| Comparison | Estimate | Std. Error | t-value | p-value |
| --- | --- | --- | --- | --- |
| Data-driven - Data-driven (bifactor) | -0.058331 | 2.611e-04 | -223.4 | <.001*** |
| RDoC - Data-driven (bifactor) | -0.132548 | 2.611e-04 | -507.6 | <.001*** |
| RDoC (bifactor) - Data-driven (bifactor) | -0.055256 | 2.611e-04 | -211.6 | <.001*** |
| RDoC - Data-driven | -0.074217 | 2.556e-04 | -290.4 | <.001*** |
| RDoC (bifactor) - Data-driven | 0.003075 | 2.556e-04 | 12.0 | <.001*** |
| RDoC (bifactor) - RDoC | 0.077292 | 2.556e-04 | 302.4 | <.001*** |

**Table for pairwise comparison of AIC Fit Measure**

| Comparison | Estimate | Std. Error | t value | p-value |
| --- | --- | --- | --- | --- |
| Data-driven - Data-driven (bifactor) | -2084.535 | 6.387 | -326.4 | <.001*** |
| RDoC - Data-driven (bifactor) | 2799.962 | 6.387 | 438.4 | <.001*** |
| RDoC (bifactor) - Data-driven (bifactor) | 1017.953 | 6.386 | 159.4 | <.001*** |
| RDoC - Data-driven | 4884.497 | 6.252 | 781.2 | <.001*** |
| RDoC (bifactor) - Data-driven | 3102.487 | 6.251 | 496.3 | <.001*** |
| RDoC (bifactor) - RDoC | -1782.010 | 6.252 | -285.0 | <.001*** |

**Table for pairwise comparison of BIC Fit Measure**

| Comparison | Estimate | Std. Error | t-value | p-value |
| --- | --- | --- | --- | --- |
| Data-driven - Data-driven (bifactor) | -2142.274 | 6.387 | -335.4 | <.001*** |
| RDoC - Data-driven (bifactor) | 2699.880 | 6.387 | 422.7 | <.001*** |
| RDoC (bifactor) - Data-driven (bifactor) | 1014.103 | 6.386 | 158.8 | <.001*** |
| RDoC - Data-driven | 4842.154 | 6.252 | 774.5 | <.001*** |
| RDoC (bifactor) - Data-driven | 3156.378 | 6.251 | 504.9 | <.001*** |
| RDoC (bifactor) - RDoC | -1685.776 | 6.252 | -269.7 | <.001*** |

**B.**

**Table for pairwise comparison of RMSEA Fit Measure**

| Comparison | Estimate | Std. Error | t-value | p-value |
| --- | --- | --- | --- | --- |
| Data-driven - Data-driven (bifactor) | 4.07E-02 | 4.26E-05 | 955.94 | <.001*** |
| RDoC - Data-driven (bifactor) | 4.36E-03 | 4.26E-05 | 102.4 | <.001*** |
| RDoC (bifactor) - Data-driven (bifactor) | 4.61E-04 | 4.26E-05 | 10.82 | <.001*** |
| RDoC - Data-driven | -3.63E-02 | 4.26E-05 | -853.54 | <.001*** |
| RDoC (bifactor) - Data-driven | -4.02E-02 | 4.26E-05 | -945.12 | <.001*** |
| RDoC (bifactor) - RDoC | -3.90E-03 | 4.26E-05 | -91.58 | <.001*** |

**Table for pairwise comparison of CFI Fit Measure**

| Comparison | Estimate | Std. Error | t-value | p-value |
| --- | --- | --- | --- | --- |
| Data-driven - Data-driven (bifactor) | -0.25038 | 0.00024 | -1043.66 | <.001*** |
| RDoC - Data-driven (bifactor) | -0.04249 | 0.00024 | -177.126 | <.001*** |
| RDoC (bifactor) - Data-driven (bifactor) | -0.00152 | 0.00024 | -6.316 | <.001*** |
| RDoC - Data-driven | 0.207887 | 0.00024 | 866.528 | <.001*** |
| RDoC (bifactor) - Data-driven | 0.248865 | 0.00024 | 1037.339 | <.001*** |
| RDoC (bifactor) - RDoC | 0.040979 | 0.00024 | 170.811 | <.001*** |

**Table for pairwise comparison of TLI Fit Measure**

| Comparison | Estimate | Std. Error | t-value | p-value |
| --- | --- | --- | --- | --- |
| Data-driven - Data-driven (bifactor) | -0.24717 | 0.000247 | -1000.3 | <.001*** |
| RDoC - Data-driven (bifactor) | -0.02489 | 0.000247 | -100.74 | <.001*** |
| RDoC (bifactor) - Data-driven (bifactor) | -0.00265 | 0.000247 | -10.71 | <.001*** |
| RDoC - Data-driven | 0.222274 | 0.000247 | 899.55 | <.001*** |
| RDoC (bifactor) - Data-driven | 0.244521 | 0.000247 | 989.59 | <.001*** |
| RDoC (bifactor) - RDoC | 0.022248 | 0.000247 | 90.04 | <.001*** |

**Table for pairwise comparison of AIC Fit Measure**

| Comparison | Estimate | Std. Error | t value | p-value |
| --- | --- | --- | --- | --- |
| Data-driven - Data-driven (bifactor) | 6767.221 | 8.538 | 792.58 | <.001*** |
| RDoC - Data-driven (bifactor) | 1111.047 | 8.538 | 130.126 | <.001*** |
| RDoC (bifactor) - Data-driven (bifactor) | 16.87 | 8.538 | 1.976 | .197 |
| RDoC - Data-driven | -5656.18 | 8.538 | -662.454 | <.001*** |
| RDoC (bifactor) - Data-driven | -6750.35 | 8.538 | -790.604 | <.001*** |
| RDoC (bifactor) - RDoC | -1094.18 | 8.538 | -128.151 | <.001*** |

**Table for pairwise comparison of BIC Fit Measure**

| Comparison | Estimate | Std. Error | t-value | p-value |
| --- | --- | --- | --- | --- |
| Data-driven - Data-driven (bifactor) | 6697.934 | 8.538 | 784.465 | <.001*** |
| RDoC - Data-driven (bifactor) | 976.32 | 8.538 | 114.347 | <.001*** |
| RDoC (bifactor) - Data-driven (bifactor) | 24.568 | 8.538 | 2.877 | .0211* |
| RDoC - Data-driven | -5721.61 | 8.538 | -670.118 | <.001*** |
| RDoC (bifactor) - Data-driven | -6673.37 | 8.538 | -781.588 | <.001*** |
| RDoC (bifactor) - RDoC | -951.752 | 8.538 | -111.47 | <.001*** |

Supplementary Table 3. Table of all RDoC seed terms used to source coordinate activation maps using Neurosynth. Maps based on these terms formed the dataset for external validation. AR: Arousal and Regulatory systems; CS: Cognitive Systems; NV: Negative Valence systems; PV: Positive Valence systems; SS: Sensorimotor Systems; SP: Social Processes.

| No. | Domain | RDoC construct | Seed terms |
| --- | --- | --- | --- |
| 1 | AR | Arousal | Arousal |
| 2 | AR | Circadian Rhythms, Sleep-wakefulness | Circadian/sleep/wakefulness |
| 3 | CS | Attention | Attention |
| 4 | CS | Cognitive Control - Goal Selection | Goal selection |
| 5 | CS | Cognitive Control - Performance Monitoring | Performance monitoring |
| 6 | CS | Cognitive Control - Response Selection | Response selection |
| 7 | CS | Cognitive Control - Inhibition/Suppression | Suppression |
| 8 | CS | Declarative Memory | Declarative memory |
| 9 | CS | Declarative Memory/Working Memory | Memory |
| 10 | CS | Language | Language |
| 11 | CS | Perception - Auditory Perception | Auditory perception |
| 12 | CS | Perception - Olfactory/Somatosensory/Multimodal/Perception | Multimodal |
| 13 | CS | Perception - Olfactory/Somatosensory/Multimodal/Perception | Olfactory perception |
| 14 | CS | Perception - Olfactory/Somatosensory/Multimodal/Perception | Somatosensory perception |
| 15 | CS | Perception - Visual Perception | Visual perception |
| 16 | CS | Working Memory - Interference Control | Interference control |
| 17 | CS | Working Memory - Active Maintenance, Limited Capacity | Working memory maintenance/capacity |
| 18 | CS | Working Memory - Flexible Updating | Working memory updating |
| 19 | NV | Acute Threat ("Fear") | Fear |
| 20 | NV | Loss | Loss |
| 21 | NV | Potential Threat ("Anxiety") | Anxiety |
| 22 | PV | Reward Learning - Reward Prediction Error | Prediction error |
| 23 | PV | Reward Learning - Probabilistic and Reinforcement Learning | Probabilistic/reinforcement learning |
| 24 | PV | Reward Responsiveness - Reward Anticipation | Reward anticipation |
| 25 | PV | Reward Responsiveness | Reward response |
| 26 | PV | Reward Valuation - Reward (probability) | Reward probability |
| 27 | SP | Affiliation and Attachment | Attachment |
| 28 | SP | Perception and Understanding of Others - Animacy Perception | Animacy |
| 29 | SP | Perception and Understanding of Others - Action Perception | Action perception |
| 30 | SP | Social Communication | Communication |
| 31 | SS | Agency and Ownership | Sensory agency/sensory ownership |
| 32 | SS | Motor Actions - Execution | Execution |
| 33 | SS | Motor Actions | Motor action |
| 34 | SS | Motor Actions - Inhibition and Termination | Motor inhibition |
| 35 | SS | Motor Actions - Initiation | Motor initiation |
| 36 | SS | Innate Motor Patterns | Motor pattern |

Supplementary Table 4: Table of spatial autocorrelation-adjusted p-values for factor score correlations shown in Figure 4B.

|  | F2 | F1 | F8 | F4 | F3 | F6 | F5 | F7 |
| --- | --- | --- | --- | --- | --- | --- | --- | --- |
| SS | <.001*** | .303 | .468 | .639 | .179 | .089 | .427 | .439 |
| SP | .545 | <.001*** | .499 | .275 | .816 | .632 | .73 | .052 |
| PV | .723 | .983 | .003** | .762 | .393 | .121 | .703 | .595 |
| NV | .807 | .962 | .489 | .432 | .884 | .939 | .63 | .202 |
| CS | .559 | .5 | .53 | .946 | .565 | .566 | .703 | .98 |
